# Relationships between biomechanical parameters, neurological recovery, and neuropathology following concussion in swine

**DOI:** 10.1101/2021.02.09.430268

**Authors:** Kathryn L. Wofford, Michael R. Grovola, Dayo O. Adewole, Kevin D. Browne, Mary E. Putt, John C. O’Donnell, D. Kacy Cullen

## Abstract

Mild traumatic brain injury (mTBI) affects millions of individuals annually primarily through falls, traffic collisions, or blunt trauma and can generate symptoms that persist for years. Closed-head rotational injury is the most common form of mTBI and is defined by a rapid change in acceleration within an intact skull. Injury kinematics – the mechanical descriptors of injury-inducing motion – explain movement of the head, energy transfer to the brain, and, therefore, determine injury severity. However, the relationship between closed-head rotational injury kinematics – such as angular velocity, angular acceleration, and injury duration – and outcome after mTBI is currently unknown. To address this gap in knowledge, we analyzed archived surgical records of 24 swine experiencing a diffuse closed-head rotational acceleration mTBI against 12 sham animals. Kinematics were contrasted against acute recovery outcomes, specifically apnea, extubation time, standing time, and recovery duration. Compared to controls, animals with mTBI were far more likely to have apnea (p<0.001) along with shorter time to extubation (p=0.023), and longer time from extubation to recovery (p=0.006). Using regression analyses with variable selection, we generated simplified linear models relating kinematics to apnea (R^2^=0.27), standing time (R^2^=0.39) and recovery duration (R^2^=0.42). Neuropathology was correlated with multiple kinematics, with maximum acceleration exhibiting the strongest correlation (R^2^=0.66). Together, these data suggest the interplay between multiple injury kinematics, including minimum velocity and middle to minimum acceleration time, best explain acute recovery parameters and neuropathology after mTBI in swine. Future experiments that independently manipulate individual kinematics could be instrumental in developing translational diagnostics for clinical mTBI.

**HIGHLIGHTS:**

1. Acute recovery parameters including apnea, extubation time, and recovery duration were altered after a single closed-head mTBI in swine.
2. Lasso-based regressions utilized kinematic parameters, including minimum velocity and middle to minimum acceleration time, to relate kinematics to apnea time, standing time, and recovery duration.
3. Lasso regression equations were able to modestly predict apnea time (R^2^=0.27) and moderately predict standing time (R^2^=0.39) and recovery duration (R^2^=0.42).
4. Injury kinematic parameters, primarily maximum acceleration, were correlated with white matter pathology after mTBI.

## INTRODUCTION

Traumatic brain injury (TBI) is a major cause of death and disability, affecting approximately 69 million individuals annually across the globe (Dewan *et al.*, 2019). Mild TBI (mTBI) comprises the vast majority of TBIs and can induce symptoms that persist for months to years including but not limited to memory loss, sleep disruptions, emotional disturbances, headaches, fatigue, nausea, sensitivity to light, sensitivity to sounds, confusion, and slowed thinking (Langlois *et al.*, 2006; Katz *et al.*, 2015). For the vast majority of people experiencing a mTBI (frequently termed concussion), symptoms progressively abate and completely resolve within weeks or months after the injury (Alves *et al.*, 1993; Levin and Diaz-Arrastia, 2015). However, the remaining 10-40% of people who received a mTBI will experience injury-induced symptoms that can persist for many months to years, affecting their ability to hold a job, emotional stability, and quality of life (Alves *et al.*, 1993; Langlois *et al.*, 2006; Mac Donald *et al.*, 2017).

Closed-head rotational injuries, the most common form of mTBI, are typically induced through rapid acceleration and deceleration of the head (LaPlaca *et al.*, 2007; Centers for Disease Control and Prevention, 2014; Helmick *et al.*, 2015; Meaney and Cullen, 2016). These movements transmit forces to the brain tissue causing diffuse damage that is difficult to detect with current clinical practices. Injury kinematics, or the mechanical description of injury motion (e.g. duration of movement, angular acceleration, angular velocity, etc.), explain movement of the head and energy transfer to the brain, which determine injury severity (Margulies and Thibault, 1992; Geddes *et al.*, 2003; Greenwald *et al.*, 2008; Rowson *et al.*, 2012). However, the relationship between specific closed-head injury kinematics – such as peak angular velocity, minimum angular acceleration, and positive acceleration duration – and outcomes after mTBI is currently unknown. In order to investigate this missing link, we explored the relationship between injury kinematics, acute recovery times, and neuropathology after a mTBI in swine. Within this study we completed a retrospective analysis of archived data that were generated with a porcine model of closed-head diffuse rotational acceleration mTBI. A swine model was utilized because it can recapitulate the mechanical loading conditions observed in clinical presentations of mTBI, such as diffuse shear deformation forces which are the primary means through which most human brain injuries are generated (Holbourn, 1943; Kleiven, 2013; Meaney and Cullen, 2016; Keating and Cullen, 2021). Furthermore, previous literature found that this injury model can simultaneously generate pathological and functional alterations that are observed in clinical TBI (Smith *et al.*, 1999, 2000; Browne *et al.*, 2011; Cullen *et al.*, 2016; Wofford *et al.*, 2017, 2019; Wolf *et al.*, 2017; Johnson *et al.*, 2018). We postulated that acute recovery parameters including apnea time, time to extubation, return to weight bearing posture, and recovery duration would increase in animals experiencing a mTBI. Furthermore, we expected that injury kinematics would be major drivers of acute recovery parameters, and as such, we expected that we could generate explanatory mathematical models relating injury kinematics to acute recovery for the first time.

## METHODS

### Animal Handling and Anesthesia

All swine procedures were completed in accordance with the Guide for the Care and Use of Laboratory Animals (US National Research Council, 2011) and followed the ARRIVE Guidelines. All protocols were approved by the University of Pennsylvania’s Animal Care and Use Committee. For the current study, surgical records and injury recordings were collected from a previously completed study of adult castrated male Yucatan miniature pigs with an average weight of 34 kg. Food and water were provided *ad libitum* and animals were housed indoors in a facility accredited by the Association for Assessment and Accreditation of Laboratory Animal Care International.

Animals were randomly assigned to either a sham procedure (n=12) or to an injury procedure (n=24). Animals were fasted overnight prior to the injury with water remaining *ad libitum*. Animals were induced with a cocktail of ketamine (12-26 mg/kg) and midazolam (0.3-0.6 mg/kg). All animals were intubated with endotracheal tube and anesthesia was maintained at 1-5% isoflurane per 2-3 liters of 100% O_2_ for the duration of the procedure. Animals were given 0.01 mg/kg glycopyrrolate subcutaneously and eye lubricant was applied. Animals receiving an injury were given 50 mg/kg acetaminophen per rectum. Physiological monitoring of heart rate, respiratory rate, and arterial oxygen saturation allowed titration of anesthesia so that all values were within acceptable ranges (heart rate between 100-130 beats per minute, respirations between 9-12 breaths per minute, and SpO_2_ between 97-100%). A forced-air temperature management system was used to maintain normothermia throughout the procedure.

Isoflurane volume was calculated as:

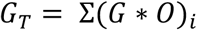

Where *G_T_* is the total isoflurane gas provided by anesthesia equipment; *G_i_* is the percentage of isoflurane gas set by the anesthetist at each time interval and *O_i_* is the rate of oxygen delivered through the anesthesia regulator at each time interval. Weighted anesthesia score was calculated as:

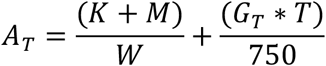

Where *K* is mass of administered ketamine; *M* is the mass of administered midazolam; *W* is the animal subject’s weight; and *T* is the amount of time the subject was under anesthesia.

### Closed-head Diffuse Brain Injury & Recovery Procedure

Rotational acceleration closed-head diffuse mTBI was completed while animals were under anesthesia by utilizing a HYGE pneumatic actuator. The animal’s mouth was positioned around a padded bite plate and then secured to the device with adjustable snout straps. The HYGE device and accompanying linkage assembly were constructed to move the head in the coronal plane (circumferential to the brainstem) in order to produce a purely impulsive non-impact head rotation (Cullen *et al.*, 2016) (**Figure 1A**). The device rapidly accelerates the swine’s head and induces forces scalable to clinical TBIs (Cullen *et al.*, 2016). Within this cohort, swine were subjected to coronal rotational injuries that ranged from to 165 to 270 radians/second to induce a mTBI. Angular displacement over time was recorded with a magneto-hydrodynamic sensor (Applied Technology Associates, Albuquerque, NM) connected to a National Instruments DAQ, controlled by LabVIEW. The sampling rate for the sensors was 10kHz. Sham animals received all other procedures absent head rotation.

**Figure 1.**
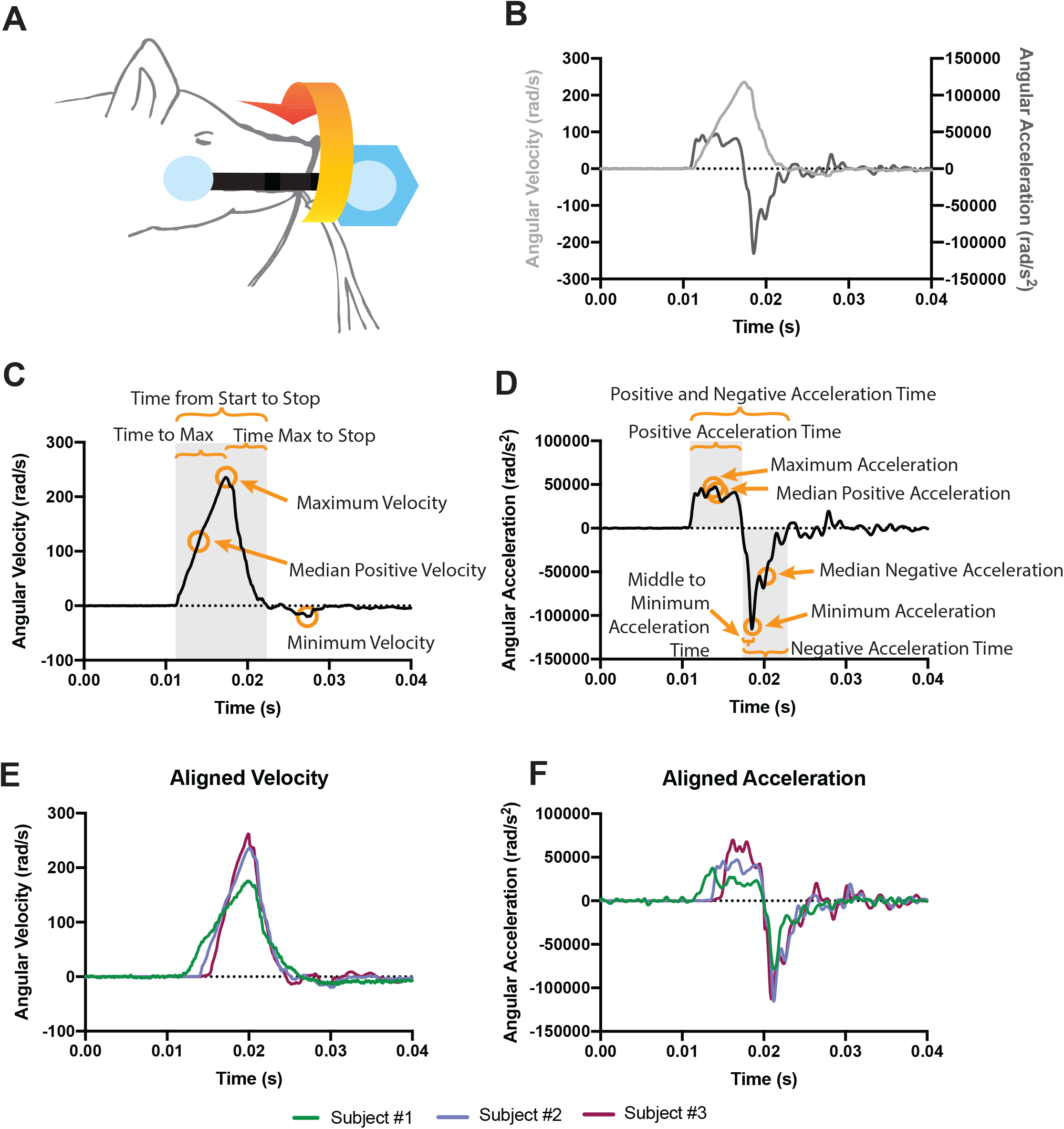
Schematic of injury model and resulting kinematic parameters. (A) Anesthetized swine assigned to the injury cohort were subjected to a rotational angular acceleration injury in the coronal plane (adapted from (Cullen *et al.*, 2016) with permission). The curved arrow indicates the rotational motion of the head during injury. (B) Movement kinematics were collected so that angular velocity and angular acceleration traces could be plotted for each injury. (C) A sample angular velocity trace and six resulting kinematic parameters describing the injury. (D) A sample acceleration trace and eight resulting kinematic parameters describing the injury. (E) Selected angular velocity and (F) angular acceleration traces from three different animal subjects exemplify differences in velocity magnitude, acceleration peaks, and acceleration durations. Subject #1 (green line) is an example of a lower maximum velocity that occurs over a longer period of time, generating a lower, but sustained positive acceleration phase and a diminished but extended negative acceleration phase. Subject #2 (blue line) is an example of a high maximum velocity that occurs over a moderate period of time, generating a higher and sustained positive acceleration phase. Subject #3 (purple line) is an example of a high maximum velocity that occurs over a short period of time, generating a high but short positive acceleration phase.

Immediately following injury, apnea was measured as the number of seconds the animal subject did not draw breath independently. Following the injury, swine were removed from the bite plate and examined for any oral or dental injuries. Once animals were returned to their housing units, isoflurane was turned off, but oxygen continued to be delivered through the endotracheal tube. Extubation time was measured as the time from isoflurane removal to the time in which animals were extubated, as prompted by chewing on the endotracheal tube, swallowing, or coughing. Standing time was measured as the time from isoflurane removal to the time in which animals were weight-bearing on all four limbs. Recovery duration was defined as the difference between standing and extubation times. Swine were continuously monitored for the duration of the recovery process. Animal data utilized in this study were from n=12 sham and n=24 mTBI archived surgical records.

### Calculating Injury Kinematics

Angular velocity of the injury was calculated by taking the derivative of the averaged position data from the magneto-hydrodynamic sensors and was smoothed with a binomial smoothing algorithm as a low-pass filter to remove chatter. Thereafter, the derivative of the smoothed velocity trace was taken to generate angular acceleration during the injury (**Figure 1B**). Six different kinematic parameters were collected from the velocity traces from each animal including: maximum velocity; median positive velocity; minimum velocity; time to maximum velocity; time from maximum velocity to stop velocity; and time from start of injury to stop velocity (**Figure 1C**). Eight different kinematic parameters were collected from the acceleration traces from each animal including: maximum acceleration; median positive acceleration; median negative acceleration; minimum acceleration; positive acceleration time; middle to minimum acceleration time; negative acceleration time; and positive and negative acceleration time (**Figure 1D**).

Jerk is defined as the rate of change in acceleration. We were most interested in measuring the rate of change between maximum acceleration and minimum acceleration, or the maximum jerk. We calculated the maximum jerk as the largest value in the vector:

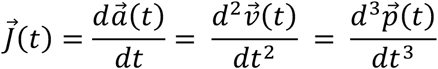

Where *a* is acceleration; *v* is velocity; *p* is position; and *t* is time. Excursion was defined as the distance traveled during the injury in radians. We calculated excursion at maximum velocity as:

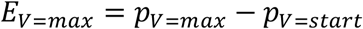

Where *p_V=max_* is position at maximum velocity and *p_V=start_* is position at the start of the injury. We calculated total excursion of the injury as:

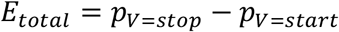

Where *p_V=stop_* is position at the end of the injury. All kinematic traces were processed and individual parameters were calculated in MATLAB version 2019b (9.7.0.1247435).

### Sacrifice and Tissue Acquisition

At the designated time point, animals were induced and intubated as described above. Thereafter, animals were transcardially perfused with 0.9% heparinized saline followed by 10% neutral buffered formalin. Decapitated tissue was stored overnight in 10% neutral buffered formalin before the brain was extracted and post-fixed in 10% neutral buffered formalin for one week at 4°C. Brains were blocked coronally every 5mm, paraffin embedded, and 8μm sections were collected via rotary microtome.

### Pathological Characterization

Assessment of diffuse axonal injury, a key characteristic of TBI pathology, was previously quantified and reported for this animal cohort (Grovola *et al.*, 2020). Briefly, the extent of amyloid precursor protein (APP) was measured by an analyst blinded to injury condition on an ordinal scale from 0 to 3 with 0 representing no pathology and 3 representing severe pathology. Because the extent of APP pathology changes over time after the injury, we only considered pathological scores from animals sacrificed 7 days after injury. Pathology scores were available for periventricular white matter, striatum, ventral thalamus, dorsal thalamus, hippocampus/fornix, and cerebellum. The total APP pathology score was generated by taking the mean pathology score across these six regions. The correlative relationships between pathology and several injury kinematics was described with a line of best fit and 95% confidence interval bands.

### Statistical Analysis

As described above, archived records from swine surgical procedures were utilized to gather information related to the dose and duration of anesthesia, as well as the time to recovery after a sham or mTBI procedure. Twenty-four injured animals and twelve sham animals meeting the inclusion criteria of having complete and interpretable surgical records were utilized in this study. As none of the sham animals exhibited apnea we categorized apnea as present or absent and carried out a Fisher’s Exact test to assess differences in the proportion of animals with apnea. To compare sham and injured animals, Welch two sample t-tests were utilized to assess mean differences in extubation time, standing time, and recovery duration. Pairwise correlations were visualized to investigate associations between outcome measures and anesthesia for all animals (**Sup. Figure 1**); between outcome measures and kinematics for all animals (**Sup. Figure 2**); and between outcome measures and kinematics for mTBI animals (**Figure 3**).

Principal component analysis (PCA) was used as a method for dimensionality reduction within the mTBI animal cohort. Several of the kinematic variables were highly collinear with one another. To stabilize the model, we used only kinematic variables with a pairwise Spearman’s correlation coefficient |*ρ|* < 0.9, leaving twelve kinematics as input variables, which were Z-scored and used for PCA. Linear models were generated for each outcome metric with the first four principal components as additive input variables. Because PC2 (the second principle component) explained the largest proportion of variance, we generated scatter plots of PC2 values against apnea time, extubation time, standing time, and recovery duration for each injured animal.

Next, we modeled the relationship between outcome metrics and key kinematic predictor variables using a least absolute shrinkage and selection operator (lasso) regression. The same 12 kinematic variables used for PCA, were used for the lasso regression. With 12 potential predictor variables but only 24 TBI animals, the lasso was necessary to reduce the number of predictor variables in the model (Tibshirani, 1996). After using the lasso to select a subset of the variables we refit the linear regression models and plotted the predicted versus the experimental values. A line of best fit with 95% confidence interval bands and the R^2^ value were reported for each relationship.

For all tests, the type I error rate was set to 0.05 and all hypothesis tests were two-sided. All statistical analyses were carried out in R Studio Version 3.6.3 using R and the *Hmisc*, *stats*, *rstatix*, *corrplot*, *Matrix*, *glmnet*, *factoextra*, and *ggplot2* packages. Line graphs, column graphs, and scatter plots were generated in Prism version 8.4.3(471).

## RESULTS

### Kinematic parameters of closed-head diffuse TBI influence recovery outcomes

Mild TBI was generated in male Yucatan mini swine with an inertial closed-head diffuse brain injury in the coronal plane. Angular velocity and angular acceleration were plotted over time for each injury and 17 resulting kinematic parameters were collected to describe the injury (**Figure 1**). Deployment of the HYGE injury device allows for control over the maximum velocity experienced by the animal subject. As a result, injured animals in this cohort intentionally experienced coronal TBIs ranging from to 165 to 270 rad/s. However, maximum velocity is not modulated in isolation. In general, when maximum velocity decreases, maximum acceleration decreases, minimum acceleration (e.g., maximum deceleration) increases, the duration of the positive acceleration phase increases, and the duration of the negative acceleration phase increases (**Figure 1E-F**). After mTBI, none of the control animals experienced apnea while 80% of the injured animals experienced apnea that ranged from 7 to 44 seconds (p<0.001; 95% CI = 0.000 - 0.146). Contrary to our expectations, mean time to extubation declined by 4.58 minutes (p = 0.023; 95% CI = 0.706 - 8.461) in injured animals relative to sham animals. The mean time to standing was extended by 12.71 minutes in injured animals although this trend was not significant (p=0.063; 95% CI = 26.169 - 0.753). The mean recovery duration was extended by 17.29 minutes in injured relative to non-injured animals (p = 0.006; 95% CI = 5.382 - 25.417) (**Figure 2**).

**Figure 2.**
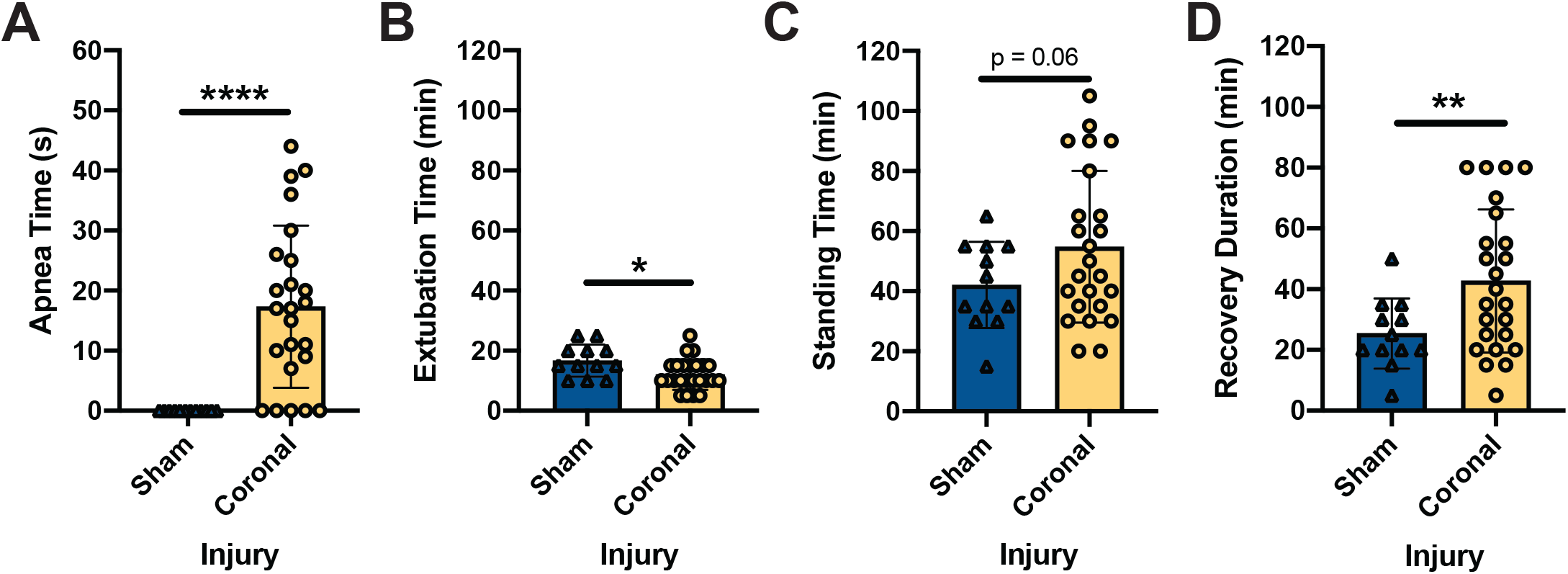
Mild TBI generated in the coronal plane alters behavioral outcome metrics acutely after injury. (A) Following mTBI, animals experienced an increase in the apnea time relative to animals receiving a sham injury (p < 0.0001). (B) The time from anesthesia removal to the time in which animals were extubated was dependent upon injury condition (p = 0.02). (C) The time from anesthesia removal to the time in which animals were weight-bearing on all four limbs was increased, but not significantly, in animals that experienced a coronal TBI (p = 0.06). (D) The time from extubation to the time in which animals were weight-bearing on all four limbs was significantly increased between sham and mTBI animals (p = 0.006). Data represent mean +/- standard deviation with * denoting p<0.05; ** denoting p<0.01; and *** denoting p<0.0001.

### Injury kinematics correlate with outcome measures

We next explored how different injury parameters related to one another using pairwise Spearman correlations. In the visual display, the upper half of the matrix depicts the correlative relationship with an ellipse while the bottom half shows the actual estimate. The shape and color of the ellipse in each cell denotes the magnitude and direction of the Spearman’s correlational coefficient (*ρ*) for the factor listed above and to the left. A correlation matrix with the four outcome parameters (black text) and the five anesthesia parameters (gray text) utilizing data from sham and injured animals appears in **Sup. Figure 1**. We did not observe strong correlations between anesthesia parameters and recovery outcomes. Ketamine dose was modestly correlated with apnea time, standing time, and recovery duration (|*ρ|* ≤ 0.36). The weighted anesthesia score, which factors in normalized ketamine, normalized midazolam, anesthesia time, and isoflurane volume was weakly correlated with each of the four outcome parameters (**Sup. Figure 1**).

Next, we generated a correlational matrix between kinematic parameters and recovery outcomes for injured animals (**Figure 3**). These data suggest the pairwise correlation between outcome measures and injury kinematics was much stronger than the correlation between outcome measures and anesthesia parameters. Among the outcome variables, apnea time was positively correlated with extubation time but was largely independent of the standing time and recovery duration. Extubation time was weakly correlated with standing time but was not correlated with recovery duaration. Standing time and recovery duration were highly correlated due to their mathematical relationship (**Figure 3**). We observed that apnea was only moderately correlated with minimum velocity and minimum acceleration (|*ρ|* ≥ 0.40) and both of these relationships were negatively correlated. Similarly, extubation time was only correlated with minimum velocity (|*ρ|* ≥ 0.40) and did not exhibit strong relationships with any other kinematic terms. In line with our finding that mTBI reduced extubation time, we observed negative correlations between extubation time and the majority of the injury kinematics. Standing time and recovery duration were correlated with many kinematic terms. Standing time had |*ρ|* ≥ 0.40 for six kinematic variables while recovery duration had |*ρ|* ≥ 0.40 for seven kinematic variables. Because recovery duration includes standing time in its calculation, it was not surprising to find that standing time and recovery duration exhibited very similar relationships to many of the kinematic terms (**Figure 3**). Time to maximum velocity, median positive acceleration, and positive acceleration time were the three most strongly correlated kinematics for standing time (|*ρ|* ≥ 0.50). Median positive acceleration and positive acceleration time were the most strongly correlated kinematics for recovery duration (|*ρ|* ≥ 0.50).

**Figure 3.**
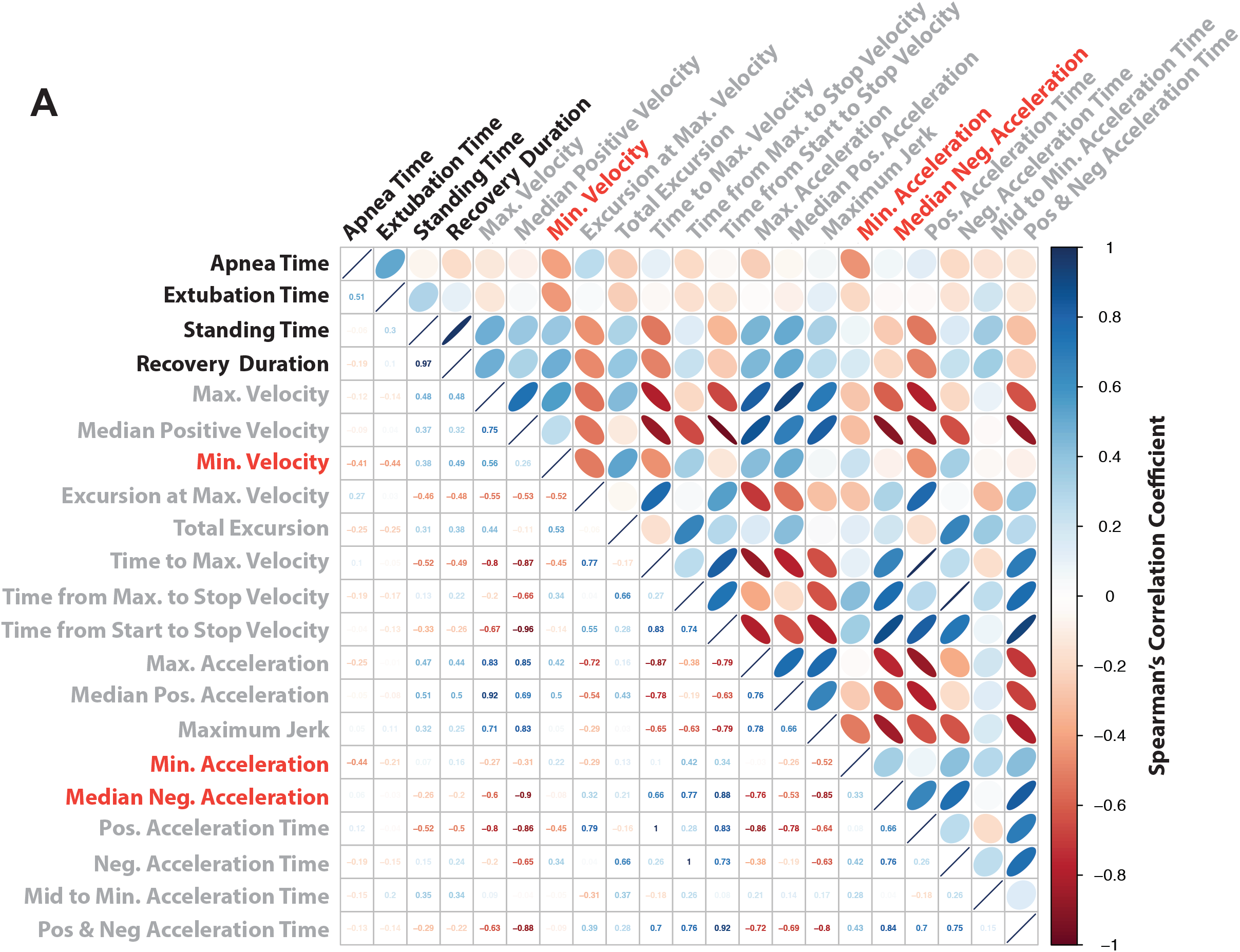
Mild TBI generated in the coronal plane alters behavioral outcome metrics acutely after injury. (A) Correlation matrix depicts the Spearman’s correlation coefficient (*ρ*) for each combination of injury parameters. Only animals experiencing an mTBI were utilized in this matrix, no sham animals were included. The matrix is organized so that the upper half of the matrix depicts the magnitude and direction of Spearman’s correlational coefficient with colored ellipses while the lower half of the matrix listed the calculated Spearman’s correlational coefficient. The orientation and color of each ellipse is indicative of the magnitude and direction of the correlation with navy ovals indicating a strong positive correlation, pastel or white circles indicating weak or no correlation, and burgundy ovals indicating a strong negative correlation. Injury parameters were colored to show outcome measures (black text), kinematic parameters with positive values (light gray text), or kinematic parameters with negative values (red text).

We generated a similar correlation matrix between outcome metrics and kinematic parameters for all sham and injured animals (**Sup. Figure 2**). Because none of the sham animals exhibited apnea, there was a point mass at the origin which artificially inflated the relationships between apnea and kinematics. Several scatter plots represented within this figure are individually plotted in **Sup. Figure 3**.

### Principal component analysis separates outcomes according to injury kinematics

Outcome metrics following mTBI were highly variable. Specifically, variability in apnea time and recovery duration suggest that factors other than just the presence or absence of injury may contribute to the spread of the data. As previously mentioned, animals within the mild coronal TBI cohort experienced vastly different injuries from one another. Indeed, maximum angular velocity ranged from 165 to 270 rad/s, maximum angular acceleration ranged from 36,004 to 126,237 rad/s^2^, and minimum angular acceleration ranged from ^−^ 66,600 to ^−^185,700 rad/s^2^. The pairwise correlations provide information about univariate associations between kinematic variables and outcome. However, we wanted to know if some combination of kinematic parameters could explain acute recovery outcomes. Therefore, we completed principal component analysis (PCA) in order to reduce the dimensionality of the mTBI animal kinematic data set. PCA was completed using 12 input kinematics that were not colinear with each other and PCA did not include outcome parameters in the analysis to ensure that any discovered relationships would not be exaggerated due to overfitting the data.

The first four principal components (PCs) had eigenvalues greater than 1, indicating that they each explained more variance than any single kinematic input parameter (**Figure 4A**). Together, the first four PCs explained 93.7% of the variance (**Figure 4B**). PC1 explained 45.9%, PC2 explained 27.6%, PC3 explained 11.6%, and PC4 explained 8.6% of the variance. Variable maps of PC1 versus PC2 and of PC3 versus PC4 illustrate the distribution and weighting of the variables across the different PCs (**Figure 4C-D**).

**Figure 4.**
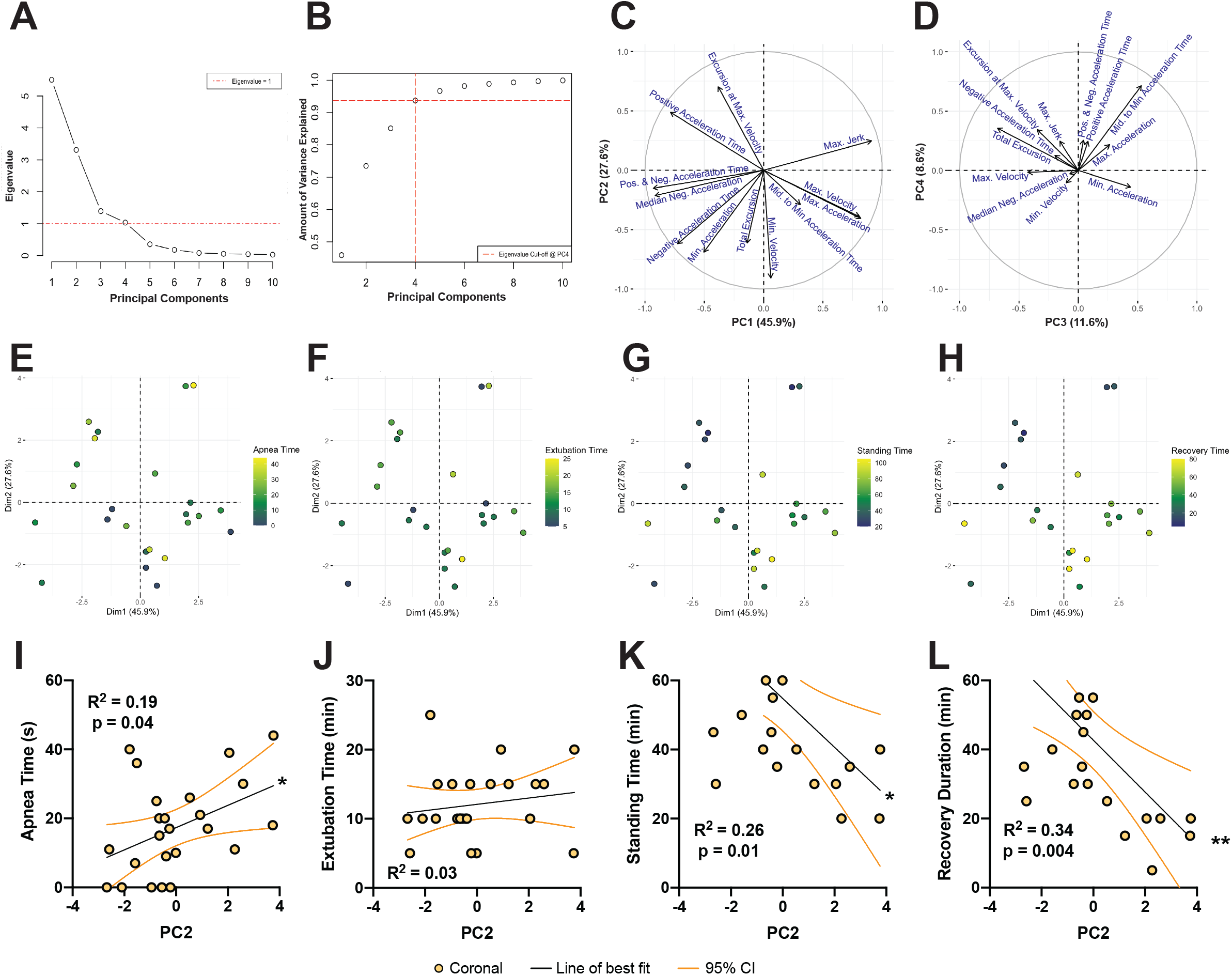
Principal component analysis is able to separate according to injury condition. PCA was completed on kinematic values from animals within the coronal cohort. (A) A scree plot and (B) a cumulative variance plot illustrate that the first four principal components had eigenvalues greater than 1 and explained 93.7% of the variance. Variance plots of (C) PC1 versus PC2 and of (D) PC3 versus PC4 show the distribution and weighting of all the input variables across each component. PC1 and PC2 were plotted against each other and were colored according to (E) apnea time, (F) extubation time, (G) standing time, and (H) recovery duration. PC1 accounted for 45.9% of the variance while PC2 accounted for 27.6% of the variance. Because PC2 best separated the outcome measures, PC2 values were plotted against (I) apnea time, (J) extubation time, (K) standing time, and (L) recovery duration. Lines of best fit (black lines) with 95% confidence intervals (orange lines) were plotted and R^2^ goodness of fit values were reported for each relationship. P values indicate PC2’s significant contributions to the outcome measure where * denotes p<0.05 and ** denotes p<0.01.

We plotted PC1 against PC2 and colored the points according to apnea time, extubation time, standing time, or recovery duration (**Figure 4E-H**). Segregation between high and low recovery outcomes was not obvious but modest separation did occur. Animals experiencing shorter apnea times generally clustered in lower values along PC2 (**Figure 4E**). PCA did not separate animals according to extubation times (**Figure 4F**). Animals experiencing shorter standing times clustered in PC2’s higher values and PC1’s lower values while animals experiencing longer standing times clustered in PC2’s lower values (**Figure 4G**). Animals experiencing shorter recovery durations clustered in PC2’s higher values and PC1’s lower values while animals experiencing longer recovery durations clustered in PC2’s lower values (**Figure 4H**).

Next, we generated linear models of the sum of the first four PCs in order to determine which PCs were most strongly related to each outcome metric (outcome ~ PC1 + PC2 + PC3 + PC4). We determined that PC2 was the only PC that was significantly associated with apnea times (p = 0.042), standing time (p = 0.014), and recovery duration (p = 0.004). None of the PCs were significantly associated with extubation time. Because PC2 values were most strongly associated with the majority of the outcomes, we plotted the PC2 value for each animal against apnea time, extubation time, standing time, and recovery duration (**Figure 4I-L**). PC2 was weakly related to apnea time (R^2^=0.19), was not at all related to extubation time (R^2^=0.03), and was modestly correlated to standing time (R^2^=0.26) and recovery duration (R^2^=0.34) (**Figure 4I-L**). The variables that most strongly contributed to PC2 were minimum velocity, excursion at maximum velocity, minimum acceleration, negative acceleration time, total excursion, and positive acceleration time.

### Injury kinematics can predict acute recovery after TBI

Our PCA results suggest that some combination of injury kinematics can explain acute recovery outcomes. Building on these findings, we used lasso regressions, a variable selection method, to identify the kinematics for a multivariable model that most strongly contributed to acute recovery outcomes. Like the PCA, we utilized the same 12 kinematic terms that were not colinear. In using the lasso regression, we determined that apnea time was best described by a liner model of three terms: minimum velocity, minimum acceleration, and middle to minimum acceleration time (**Figure 5A**). Standing time and recovery duration were both best described by linear models with six terms: maximum velocity, minimum velocity, excursion at maximum velocity, positive acceleration time, negative acceleration time, and middle to minimum acceleration time (**Figure 5A**). Two kinematic terms – minimum velocity and middle to minimum acceleration time – were preserved across the apnea time, standing time, and recovery duration models. Interestingly, the lasso regression eliminated all the input kinmetic variables when generating the most predictive model for extubation time, suggesting that a constant was better at predicting extubation time than any combination of kinematic terms.

**Figure 5.**
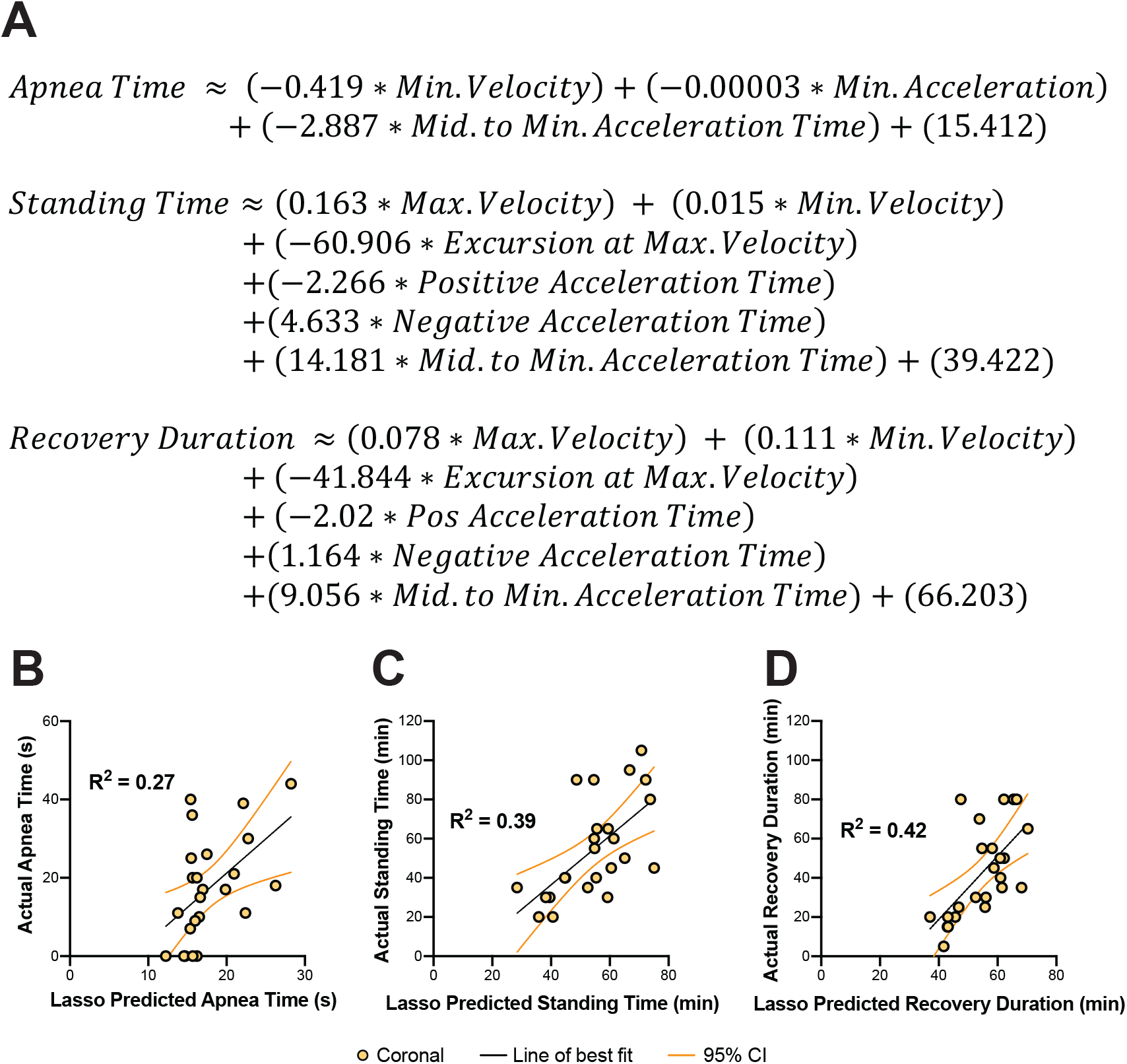
Explaining the relationships between injury kinematics and outcome measures as determined by a lasso regression. (A) Linear equations were generated to explain the relationships between injury kinematics and apnea time, standing time, and recovery duration. Thereafter, the equations were utilized to contrast (B) the predicted apnea time against the actual apnea time, (C) the predicted standing time against actual standing time, and (D) the predicted recovery duration against actual recovery duration. Lines of best fit (black lines) with 95% confidence intervals (orange lines) were plotted and R^2^ goodness of fit values were reported for each relationship.

To understand the prediction accuracy of these three equations, we next calculated the predicted apnea time, predicted standing time, and predicted recovery duration for all injured animals by plugging in each animal’s kinematic values into each equation. These predicted values were plotted against the actual, experimental outcome values. The lasso predicted values for apnea exhibited a modest correlation with the actual apnea time (R^2^=0.27; **Figure 5B**). The lasso predicted values for standing time exhibited a moderate positive correlation with the actual standing time (R^2^=0.39; **Figure 5C**). The lasso predicted values for recovery duration also exhibited a moderate positive correlation with the actual recovery duration (R^2^=0.42; **Figure 5D**).

The lasso regression was repeated on z-scored kinematic data in order to generated coefficients that could be compared against each other. For apnea, minimum velocity showed the strongest association with outcome and middle to minimum acceleration time showed the second strongest association with outcome (**Sup. Table 1**). For both standing time and recovery duration models, maximum velocity contributed most strongly to the outcomes although it seems that many of the factors contributed to outcome (**Sup. Table 1**).

### Brain pathology is correlated to injury kinematic parameters

Amyloid precursor protein (APP) accumulation in axons is the gold-standard used to identify diffuse axonal pathology following closed-head TBI (Johnson *et al.*, 2010, 2013; Tang-Schomer *et al.*, 2012; Lafrenaye, 2016). Previous efforts in our lab have characterized the distribution of APP brain pathology in these animals over time (Grovola *et al.*, 2020). However, these pathology metrics have not been compared against injury kinematics. We collected pathology data from animals surviving seven days after a sham or coronal injury. Average APP pathology was defined as the mean pathology score of 0 (no pathology) to 3 (severe pathology) across six brain regions: periventricular white matter, striatum, ventral thalamus, dorsal thalamus, fimbria/fornix, and cerebellum. We plotted the average APP pathology score against eight kinematic parameters that were important in describing the injury. We plotted average APP pathology against terms that described the magnitude of the injury: maximum velocity, maximum acceleration, minimum acceleration, and maximum jerk. We also plotted average APP against terms that were deemed important in the lasso and PCA analyses: minimum velocity, middle to minimum acceleration time, median negative acceleration, and excursion at maximum velocity.

We observed that the average APP pathology score was correlated (R^2^ > 0.45) with maximum velocity, maximum acceleration, minimum acceleration, maximum jerk, middle to minimum acceleration time, and median negative acceleration (**Figure 6A-F**). The average APP pathology score was moderately correlated with excursion at maximum velocity and had correlation with minimum velocity (**Figure 6G-H**). Maximum acceleration was the most strongly correlated with average APP pathology scores with R^2^ = 0.66.

**Figure 6.**
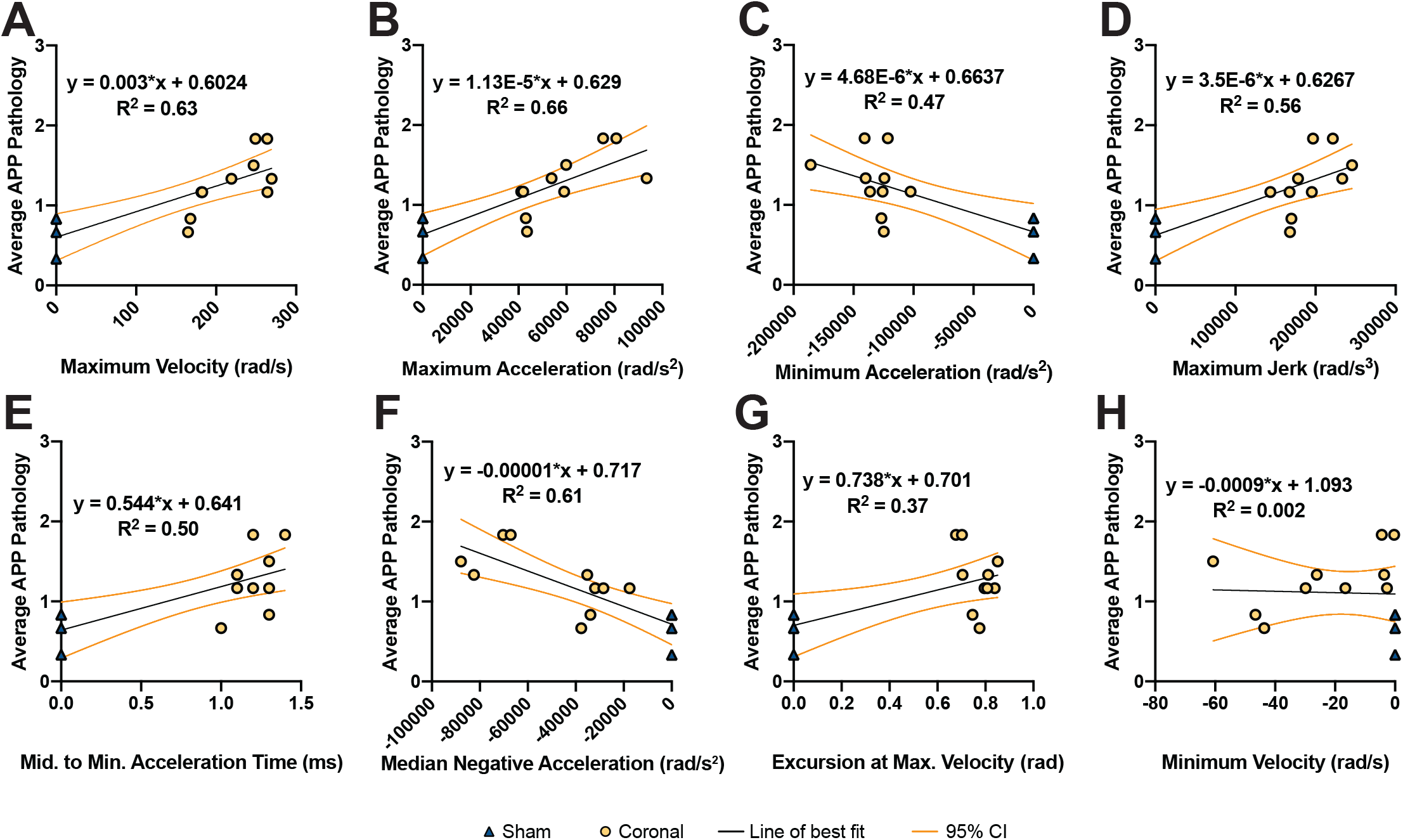
Injury kinematics are correlated with pathological burden 7 days after mTBI. Average APP burden was plotted against (A) maximum velocity, (B) maximum acceleration, (C) minimum acceleration, (D) maximum jerk, (E) middle to minimum acceleration time, (F) median negative acceleration, (G) excursion at maximum velocity, and (H) minimum velocity. Lines of best fit (black lines) and 95% confidence intervals (orange lines) were calculated for each graph and linear equations are reported for each graph. The R^2^ goodness of fit for each relationship is denoted on each graph.

We also investigated how average APP pathology correlated to our four recovery parameters: apnea time, extubation time, standing time, and recovery duration. Correlations between pathology at 7 days and apnea time, extubation time, and standing time were weakly associated, as demonstrated by R^2^ values less than 0.30 in all cases (**Sup. Figure 4A-C**). Pathology at 7 days and recovery duration was moderately correlated with an R^2^ value of 0.40 (**Sup. Figure 4D**).

## DISCUSSSION

Diffuse closed-head rotational acceleration is the most common form of TBI(LaPlaca *et al.*, 2007; Centers for Disease Control and Prevention, 2014; Helmick *et al.*, 2015; Meaney and Cullen, 2016), but the specific injury kinematic parameters of head rotation that drive pathology and affect neurological recovery are currently unknown. This retrospective study shows that acute recovery parameters including apnea time, extubation time, and recovery duration are altered after a single mTBI in swine and that a number of kinematic parameters are associated with recovery. We also found that injury kinematic parameters of closed-head diffuse injury are correlated with neuropathological outcomes after mTBI. To our knowledge, this study represents the first attempt to understand the contributions of distinct injury kinematics to recovery after injury using a large animal model of TBI. Taken together, these data inform how injury kinematic parameters affect acute outcomes and suggest that understanding the interplay between multiple injury kinematics may be key in predicting recovery following trauma.

A major goal of this work was to determine if combinations of injury kinematics could better relate to recovery outcomes. The first step in investigating the multi-component relationships was to complete PCA. We observed that of all the principal components, PC2 was most strongly associated with the recovery outcomes apnea, standing time, and recovery duration. PC2 factored in all 12 of the input kinematics but minimum velocity, excursion at maximum velocity, minimum acceleration, negative acceleration time, total excursion, and positive acceleration time were the factors that most strongly affected PC2. PCA data suggest that recovery outcomes could be better explained with a combination of kinematics instead of any singular term.

To build on this PCA finding and to improve the interpretability of the mathematical relationships to recovery parameters, we employed a lasso regression, a variable selection method. In the lasso regression analysis, we determined that minimum velocity, minimum acceleration, and middle to minimum acceleration time were the most important factors in explaining apnea time. While all three kinematics contribute to the model’s accuracy, minimum velocity was the most strongly weighted factor. This sparse linear model is consistent with the observation that apnea time was only strongly correlated with a few kinematic terms in the correlation matrix. In contrast to the apnea equation, maximum velocity, minimum velocity, excursion at maximum velocity, positive acceleration time, negative acceleration time, and middle to minimum acceleration time were the most important factors in explaining both standing time and recovery duration. These models preserved the same six kinematic inputs, which is probably due to the mathematical relationship that standing time and recovery duration share. Within these equations, maximum velocity was the most strongly weighted factor and minimum velocity was the weakest weighted factor. Minimum velocity and middle to minimum acceleration time were the only terms that were preserved in all three equations suggesting that these kinematic variables play a leading role in determining several acute recovery outcomes.

We were surprised to find that minimum velocity played a leading role in the PCA analysis and all of the lasso equations. Indeed, minimum velocity is a kinematic term that does not have a large magnitude and is not associated with the largest changes in acceleration. The minimum velocity term represents a brief change in direction of the HYGE’s linkage arm’s movement at the end of the injury, resulting in a small bounce of the animal’s head. That this small change in direction could be one of the largest drivers of acute recovery outcomes speaks to the complexity of viscoelastic tissue movement during injury. Future work controlling the magnitude and duration of the minimum velocity bounce would help better understand it’s contribution to outcomes.

We also reported correlational relationships between injury kinematics and white matter pathology 7 days after the injury. Unlike the PCA and lasso analyses, maximum acceleration, maximum velocity, and median negative acceleration had the strongest relationships with pathological outcome. Minimum velocity, which played a leading role in recovery terms, had no correlation to pathology. Middle to minimum acceleration time, another lead kinematic in explaining recovery, had a moderate correlation to pathology, although it was weaker than several other kinematics. This may suggest that the kinematics that drive acute recovery parameters are not necessarily the same kinematics that contribute to white matter pathology. This also suggests that studies utilizing closed-head rotational models should report both angular velocity kinematics, angular acceleration kinematics, and injury timing information in order for others to replicate their work. Future studies investigating why white matter pathology at 7 days did not directly relate to acute recovery parameters will also be helpful in contextualizing these findings.

Implications for developing explanatory relationships between injury kinematics and recovery outcomes are immense. Injury kinematic data can be utilized to develop computational models of brain movement during injury (Post *et al.*, 2014; Atlan *et al.*, 2018; Hajiaghamemar *et al.*, 2020). Many kinematic parameters are related to one another and computational modeling may be able to enhance predictive accuracy by informing which parameters to prioritize. If head kinematics could be measured as the injury occurs (through mounted sensors in civilian cars or on military/athlete helmets), then physicians may be able to utilize the kinematic parameters in a computational model to understand the injury severity and predict the patient’s recovery experience (Rowson *et al.*, 2012; Cortes *et al.*, 2017; Graci *et al.*, 2019; Sanchez *et al.*, 2019; Holt *et al.*, 2020; Huber *et al.*, 2021). Predictive models, like this study, could be especially useful for mTBI because no overt bleeding or pathology may be evident even though many patients report persistent effects for months or years after an injury (Alves *et al.*, 1993; Langlois *et al.*, 2006; Katz *et al.*, 2015).

This study was based on a retrospective analysis of archived injury kinematics and surgical records from swine experiencing either a sham or a mTBI. As is the nature of retrospective studies, we only utilized outcome measures that were consistently recorded across all animals. However, future studies correlating injury kinematics to long-term behavioral and physiological outcomes could be particularly informative to clinical translation of these models. Indeed, we suspect that independently modulating injury kinematics could have drastically different outcomes on pathology, physiology, and behavior (Post *et al.*, 2014; Meaney and Cullen, 2016). Because the brain is viscoelastic, the duration over which kinematics are applied to the brain determines how deep injury forces penetrate into the tissue, with rapid injuries concentrating forces on the surface of the brian and with extended injuries distributing forces into deep structures of the brain. Therefore, we suspect that longer injury durations may generate more neuronal damage, synapse disruption, and gliosis in deeper brain structures, affecting consciousness, mood, addiction, and attention (Levin *et al.*, 1988; Volkow *et al.*, 2003; Mcallister, 2011; Meaney and Cullen, 2016). In contrast, we speculate that injuries with higher maximum velocity and maximum jerk may generate more neuronal damage, synapse disruption, and gliosis in cortical brain structures, affecting aggression, memory, and coordination (Siever, 2008; Mcallister, 2011). These results illustrate that there is a need for future studies to independently control each kinematic parameter during TBI to fully understand their contributions to pathology, physiology, and behavior.

Additionally, as inertial models of closed-head diffuse TBI are becoming more common in the neurotrauma field, there is a need to compare and contrast results across institutions and investigators (Hajiaghamemar and Margulies, 2020; Mayer *et al.*, 2020). Indeed, several research groups employ a similar porcine injury model of diffuse closed-head mTBI because the architecture of swine’s large gyrencephalic brains allows loading conditions that are scalable to clinical TBIs (Johnson *et al.*, 2018; Vink, 2018; Kinder *et al.*, 2019; Hajiaghamemar and Margulies, 2020; Mayer *et al.*, 2020). Our findings suggest that maximum velocity alone may not be predictive of recovery parameters or neuropathology. Systematically controlling and reporting kinematic parameters that influence pathology and outcome will be critical to compare and contrast findings between studies.

In completing this study, we expected that an injury would delay all recovery outcome parameters. However, in contrast to our expectations, we found extubation times were significantly decreased in injured animals relative to sham animals. Future studies investigating the causal relationship between injury and extubation time are necessary to tease apart this unexpected relationship. Moreover, coupling recovery outcomes of apnea time, standing time, and recovery duration with physiological changes in the brain could be informative in interpreting these data. Indeed, investigating how pain, stress, and brainstem pathology affect extubation and standing times could support this work.

Building on this, apnea time is not commonly reported in closed-head rotation models of swine TBI. Investigating potential causes of apnea and the implications of apnea (transient reduction in oxygenation) for other behaviors after injury are important in contextualizing these findings. Furthermore, contrary to our expectations, we found that anesthesia factors – within the ranges administered in this study – did not play a major role in the outcome parameters.

While the results of this study suggest that kinematic parameters can be used in combination to predict apena time, standing time, and recovery duration; this study has several notable limitations. Most importantly, this retrospective study did not allow independent modulation of individual injury kinematics. Here, mathematical analyses attempt to weight individual kinematic parameters in order to explain outcomes. However, independently controlling one kinematic parameter at a time will facilitate a better understanding into how that type of injury affects pathology, physiology, and animal behavior. Additionally, explanatory models in this study utilized linear models with additive relationships. Modeling outcomes with non-linear relationships or investigating interactions of kinematics could enhance the prediction accuracy of these models but requires hundreds of animal replicates to be a statistically valid method. In addition to the mathematical limitations of this study, the retrospective nature of this study underscores the importance of consistent record keeping related to post-injury acute recovery metrics, thus mitigating challenges in interpreting archived procedure records. Animal recovery progress was described approximately every 10 minutes until an hour after swine were completely ambulatory. Only complete records that had descript narratives of acute recovery were included in the study. Because these acute recovery parameters were only deemed a dependent variable after all surgeries had been completed, we suspect that inaccuracies and variability would be conserved across sham and injured animals. Furthermore, personnel involved in the study would have been unaware that acute recovery times would be a part of this study and so would have acted in an unbiased manner. In this way, the retrospective study was limited in temporal resolution and accuracy, but also had the benefit of decreased observer bias. Future studies in which kinematic variables are independently modulated and dependent variables are systematically recorded could be informative in validating these results.

In addition to limitations of the study design, we were only able to correlate injury kinematics with acute metrics of recovery. However, many other parameters could be systematically quantified in swine after injury. Future swine TBI studies could record other acute outcomes including extent of vocalization, control of head movements, supporting weight on two limbs, return of coordinated walking, gait rigidity, rearing, eye movements, and oculocephalic reflexes (Datzmann *et al.*, 2019). Furthermore, the development of long-term behavioral metrics for swine to test learning, memory, aggression, anxiety, saccades, sleep disturbances, and addiction could act as a critical link in understanding injury side-effects in clinical presentations. In addition to investigating how these injuries affect behavioral outcomes, altering the plane of rotation would be critical for translation to clinical predictive models. Typically, clinical TBIs are not constrained to closed-head rotational acceleration within one plane, but rather movement of the head occurs in multiple planes during an injury. Investigating how kinematics affect recovery in injuries generated in the sagittal plane, axial plane, or oblique rotations would be important to combine with this work to predict clinical recovery. Finally, sex was not accounted for in this study, but it would be interesting to see if these trends were preserved across both male and female swine.

In summary, we utilized a large-animal model of closed-head rotational TBI that can scale loading conditions to TBI in humans. We found that acute recovery after mild closed-head diffuse TBI in swine could be modestly predicted with an additive linear model model informed by injury kinematics. Together, these data suggest that no singular kinematic parameter was predictive of outcome, rather, interplay between multiple injury kinematics drives recovery parameters and neuropathology after mTBI in swine. We believe that better understanding how injury kinematics affect neurological recovery and neuropathology will inform the development of more translational diagnostic criteria for TBI in the future.

## ACKNOWLEDGEMENTS

This work was made possible through financial support provided by the Department of Veterans Affairs [BLR&D Merit Review I01-BX003748 (Cullen); Merit Review Grant I01-RX001097 (Cullen); RR&D Career Development Award IK2-RX003376 (O’Donnell)] and the National Institutes of Health [NINDS #T32-NS043126 (Wofford); NINDS #F32-NS116205 (Wofford)]. None of the funding sources aided in the collection of the data.

## AVAILABILITY OF DATA AND MATERIALS

The datasets and computational models used within the current study are available from the corresponding author upon reasonable request.

## AUTHORS’ CONTRIBUTIONS

KLW, JCO, DKC designed the experiments; KLW, MRG, DOA, KDB, MEP performed research and analyzed data; and KLW wrote the manuscript. All authors read and approved the final manuscript.

## DIVERSITY STATEMENT

Because paper citations are considered a metric of scientific impact and because recent work found that citation practices in neuroscience fields can disproportionally cite papers authored by men in first and last author positions (Dworkin *et al.*, 2020), we strove to balance citations within this paper across man (M) and woman (W) first and last authors in proportion to available manuscripts in the neurotrauma discipline. After excluding citations of authors included in this manuscript, 49% were authored by MM (man first author and man last author), 19% were WM, 5% were MW, and 27% were WW. These citation rates are approximately proportional to the available citable papers in this field.

---

**Supplemental Figure 1. Spearman’s Rho correlation for each combination of anesthesia and recovery parameters within the sham and injured animals**. Injury parameters were colored to show outcome measures (black text) or anesthesia parameters (light gray text).

**Supplemental Figure 2. Recovery parameters across sham and injured animals are correlated with injury kinematics.**(A) Correlation matrix depicting the Spearman’s Rho correlation for each combination of kinematic and recovery parameters within the sham and injured animals. Injury parameters were colored to show outcome measures (black text), kinematic parameters with positive values (light gray text), or kinematic parameters with negative values (red text).

**Supplemental Figure 3. Injury kinematics are correlated with recovery parameters.** (A) Apnea time, (B) standing time, and (C) recovery duration were plotted against maximum velocity, maximum acceleration, minimum acceleration, maximum jerk, positive acceleration time, and negative acceleration time. Lines of best fit (solid black lines) and 95% confidence intervals (orange lines) were calculated for each graph.

**Supplemental Figure 4. White matter neuropathology is moderately correlated with recovery duration.** Average APP burden was plotted against (A) apnea time, (B) extubation time, (C) standing time, and (D) recovery duration. Lines of best fit (solid black lines) and 95% confidence intervals (orange lines) were calculated for each graph and linear equations are reported for each graph. The R^2^ goodness of fit for each relationship is denoted on each graph. Only recovery duration exhibited a moderate correlation with averaged APP pathology.

