## Supplemental Data for "Relationships between biomechanical parameters, neurological recovery, and neuropathology following concussion in swine"

### Sup Figure 1

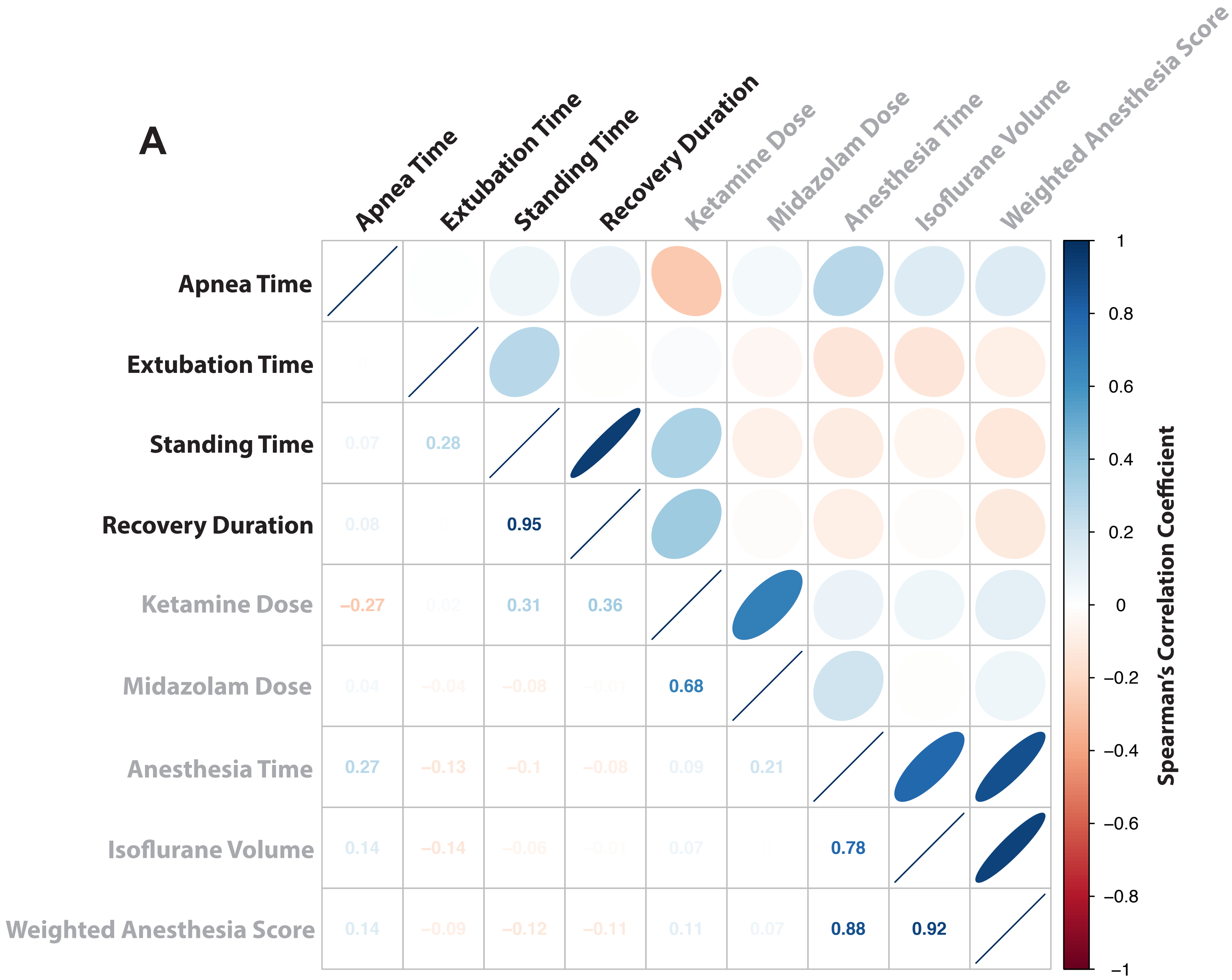


**Supplemental Figure 1. Recovery parameters are weakly correlated with anesthesia metrics.** (A) Correlation matrix depicting the Spearman's Rho correlation for each combination of anesthesia and recovery parameters within the sham and injured animals. Injury parameters were colored to show outcome measures (black text) or anesthesia parameters (light gray text).

### Sup Figure 2

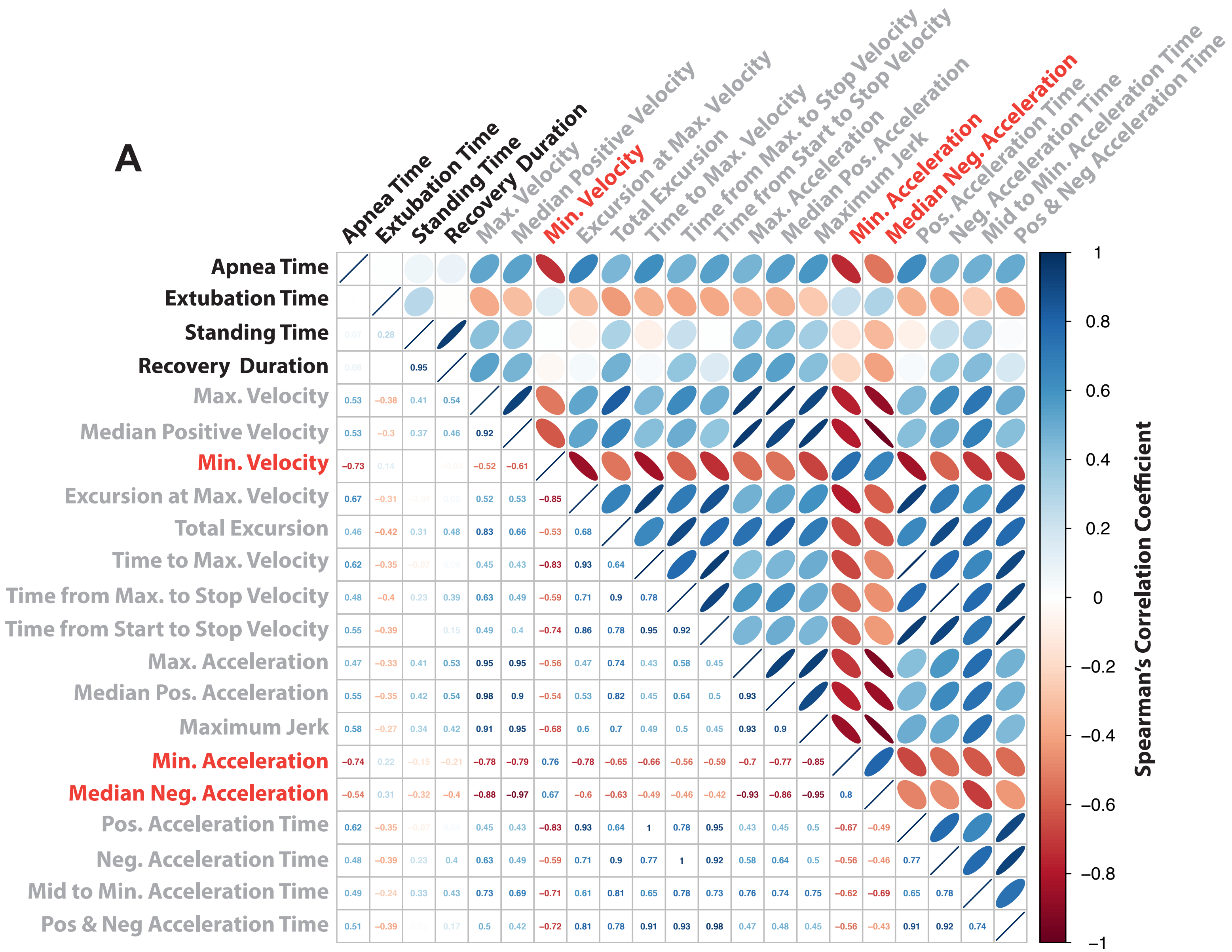


**Supplemental Figure 2. Recovery parameters across sham and injured animals are correlated with injury kinematics.** (A) Correlation matrix depicting the Spearman's Rho correlation for each combination of kinematic and recovery parameters within the sham and injured animals. Injury parameters were colored to show outcome measures (black text), kinematic parameters with positive values (light gray text), or kinematic parameters with negative values (red text).

### Sup Figure 3

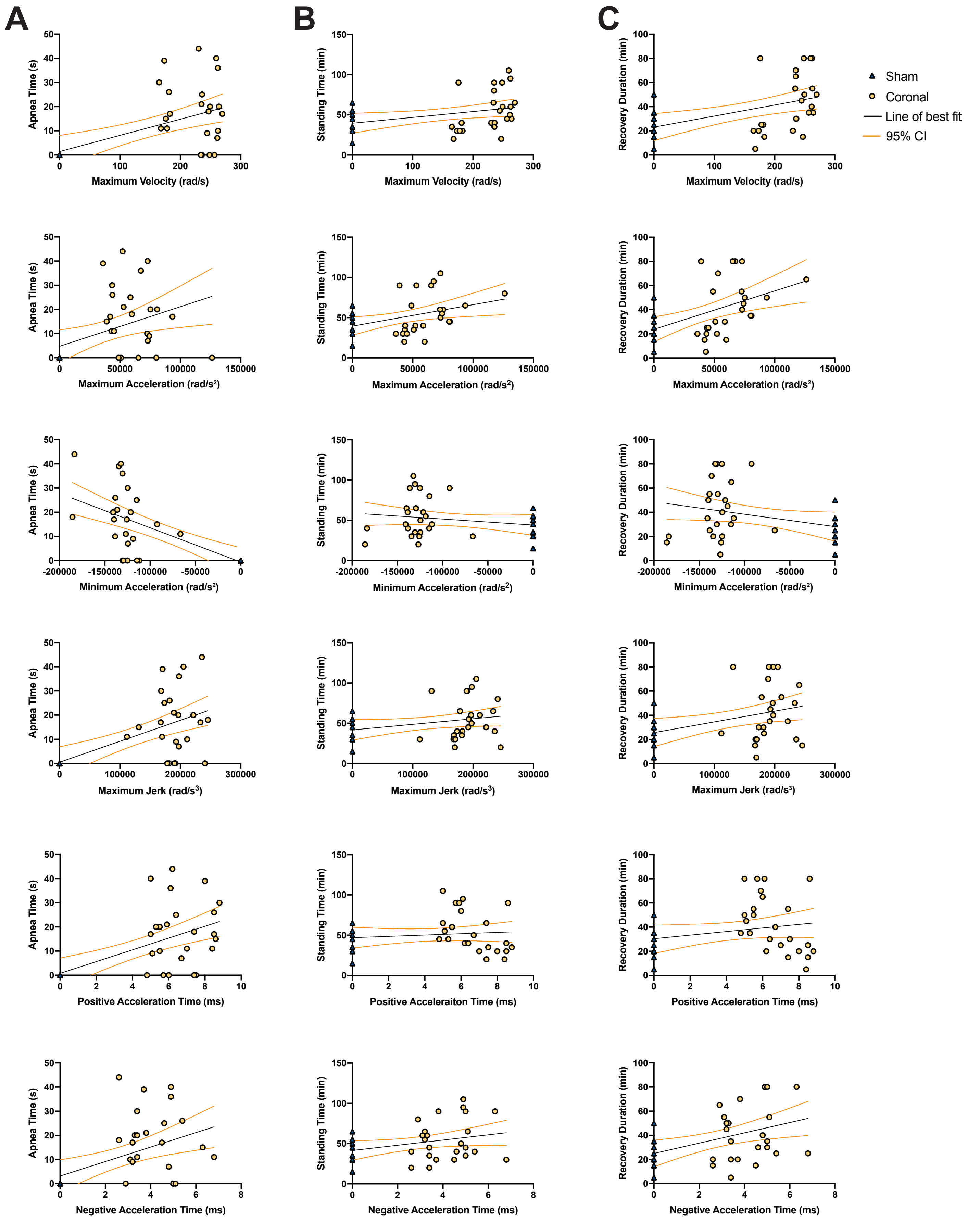


**Supplemental Figure 3. Injury kinematics are correlated with recovery parameters.** (A) Apnea time, (B) standing time, and (C) recovery duration were plotted against maximum velocity, maximum acceleration, minimum acceleration, maximum jerk, positive acceleration time, and negative acceleration time. Lines of best fit (solid black lines) and 95% confidence intervals (orange lines) were calculated for each graph.

### Sup Table 1

**Supplemental Table 1. Multivariable linear models based on variable selection using lasso methods**. Values are lasso coefficients for the raw data or for the z-scored data.

| | Coefficients for Raw Data | | | | Coefficients for Z-scored Data | | |
|---|---|---|---|---|---|---|---|
| | Apnea Time | Standing Time | Recovery Duration | | Apnea Time | Standing Time | Recovery Duration |
| Intercept | 15.412 | 39.422 | 66.203 | | 7.6E-17 | -1.3E-16 | -1.8E-16 |
| **Max. Velocity** | | 0.163 | 0.078 | | | 0.233 | 0.342 |
| **Min. Velocity** | -0.149 | 0.015 | 0.111 | | -0.278 | 0.013 | 0.100 |
| **Excursion at Max. Vel** | | -60.906 | -41.844 | | | -0.112 | -0.123 |
| **Total Excursion** | | | | | | | |
| **Max. Acceleration** | | | | | | | |
| **Max. Jerk** | | | | | | | |
| **Min. Acceleration** | -0.00003 | | | | -0.076 | | |
| **Median Neg. Acceleration** | | | | | | | |
| **Pos. Acceleration Time** | | -2.266 | -2.020 | | | -0.119 | -0.017 |
| **Neg. Acceleration Time** | | 4.633 | 1.164 | | | 0.210 | 0.240 |
| **Mid. To Min. Acceleration Time** | -2.887 | 14.181 | 9.056 | | -0.121 | 0.191 | 0.191 |
| **Pos. & Neg. Acceleration Time** | | | | | | | |

#### SUPPLEMENTAL METHODS

##### *Description of Lasso*

Given a set of possible predictor variables in a regression model, the least absolute shrinkage and selection operator (lasso) seeks to, with high probability, reduce the model to those predictors with relevant associations with outcome[1]. The method uses penalized likelihood with larger penalties yielding models with smaller numbers of predictors. The optimal penalty for any application is unknown, and typically, as in our analysis, the penalty is chosen using cross-validation to optimize prediction of the outcome. Optimal prediction often includes variables with weaker associations with outcome, and the results of a lasso applied in this way can be viewed as a screening tool. In other words, the estimated set of variables based on the lasso will with high probability, include those variables with strong associations with the outcome as well as other variables with weaker associations. Notably none of the variables are chosen using hypothesis testing, or p-values, as is typical of older algorithms such as stepwise[2].


1. Bühlmann P, Geer S van de. *Statistics for High-Dimensional Data: Methods, Theory and Applications*. Berlin Heidelberg: Springer-Verlag. Epub ahead of print 2011. DOI: 10.1007/978-3-642-20192-9.

2. Desboulets LDD. A Review on Variable Selection in Regression Analysis. *Econometrics* 2018; 6: 45.

### Sup Figure 4

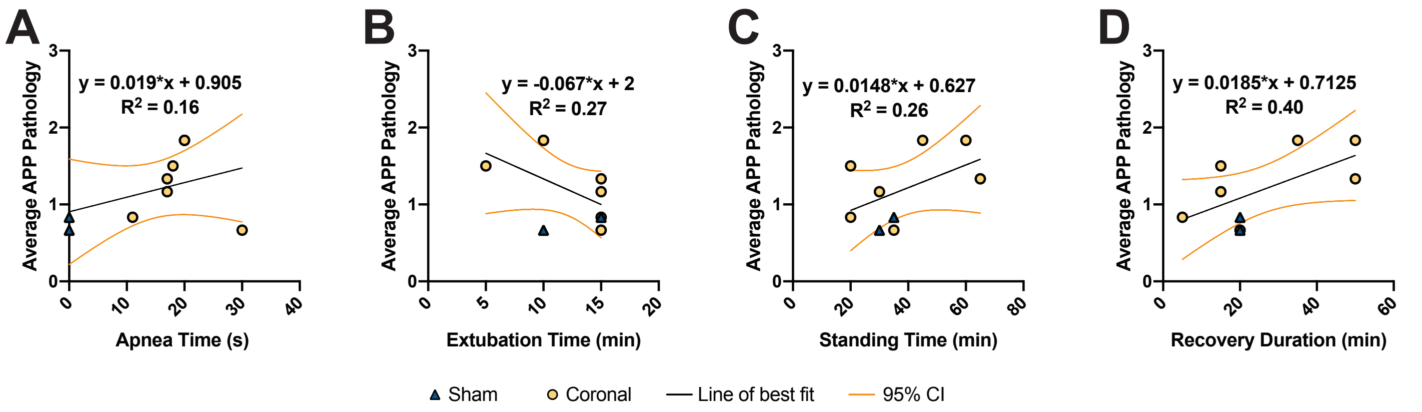


**Supplemental Figure 4. White matter neuropathology is moderately correlated with recovery duration.** Average APP burden was plotted against (A) apnea time, (B) extubation time, (C) standing time, and (D) recovery duration. Lines of best fit (solid black lines) and 95% confidence intervals (orange lines) were calculated for each graph and linear equations are reported for each graph. The R2 goodness of fit for each relationship is denoted on each graph. Only recovery duration exhibited a moderate correlation with averaged APP pathology.
